# Deciphering the genetic and transcriptional basis of cross-stress responses in *Escherichia coli* under complex evolutionary scenarios

**DOI:** 10.1101/010595

**Authors:** Violeta Zorraquino-Salvo, Semarhy Quinones-Soto, Minseung Kim, Navneet Rai, Ilias Tagkopoulos

## Abstract

A tantalizing question in microbial physiology is the inter-dependence and evolutionary potential of cellular stress response across multiple environmental dimensions. To address this question, we comprehensively characterized the cross-stress behavior of wild-type and evolved Escherichia coli populations in five abiotic stresses (n-butanol, osmotic, alkaline, acidic, and oxidative) by performing genome-scale genetic, transcriptional and growth profiling, thus identifying 18 cases of cross-stress protection and one case of cross-stress vulnerability. We identified 18 cases of cross-stress protection and one case of cross-stress vulnerability, along with core and stress-specific networks. We tested several hypotheses regarding the effect of the stress order, stress combinations, mutation reversal and varying environments to the evolution and final cellular fitness of the respective populations. Our results argue of a common systems-level core of the stress response with several crucial implicated pathways that include metal ion binding and glycolysis/gluconeogenesis that is further complemented by a stress-specific expression program.

## Introduction

Regardless of their complexity, organisms have developed a rich molecular and behavioral repertoire to cope with environmental variations in order to maintain homeostasis and cellular function. The term *stress* is used to describe conditions where environmental parameters differ substantially from an organism’s optimal conditions for growth and for bacteria it has been an active area of research for decades ^1^, given its industrial and medical importance. Most studies have focused on responses to single stressors, such as pH ^2-4^, temperature ^5-7^, oxidation ^8,9^ and UV ^10,11^, thus providing an important insight on stress-related cellular responses and their underlying mechanisms. More recently, several studies have investigated cases of stress combinations and particularly of *cross-stress protection*, where exposure in a given stressor confers a fitness advantage against an exposure to a second stress ^12-23^.

Evidence of cross-stress behavior has been mostly circumstantial so far and documented throughout the microbial kingdom. Early work on glucose- and nitrogen-starved *Escherichia coli* cells showed increase survival rates after heat shock or hydrogen peroxide-mediated stress when compared to non-stressed cells ^24^, with the alternative sigma factor *rpoH* a crucial link for heat shock protein production during starvation stress ^25^. *E. coli* cells adapted to high ethanol concentrations had decreased growth under acidic stress ^26^ and high temperature environments induce a similar transcriptional program to that observed under low oxygen ^27^. Pre-adaptation to elevated temperature can reduce both cell death rate and mutation frequency caused by hydrogen peroxide in *Lactobacillus plantarum* ^28^. Osmotic stress was found to confer inducible heat tolerance in *Salmonella typhimurium* ^15^, while trehalose synthesis, which is important for osmoprotection, was shown to feature a heat-inducible component that increases heat tolerance of osmo-adapted *Salmonella enterica* cells ^14^. More recently, gene-deletion libraries were used to investigate hydrogen peroxide (H_2_O_2_) tolerance in yeast^29^.

The emergence of cross-stress behavior and the impact of the environmental correlation-structure during its evolution remain tantalizing questions that need to be further investigated. In a recent study, the fitness of *E. coli* populations in five stressors was measured after adaptation for 500 generations under various other stressors and cross-stress dependencies were found to be ubiquitous, highly interconnected and emerging within short timeframes ^30,26,28^. While studies like that provide valuable insights on the evolutionary potential of cross-stress behavior, our knowledge of static cross-stress dependencies as well as the effect of specific environmental characteristics to their dynamic, evolutionary potential remains limited. Environmental structure is known to be important for the trait evolution during periodic fluctuations ^31-33^, co-evolution in microbial communities ^34^ and can influence the rate and direction of evolution. Additionally, cross-stress responses may be dependent on the type of co-evolution or the order of exposure to the pair of stresses, while it is not clear if and in which cases, evolved mutants revert to wild-type phenotypes in the prolonged absence of the stressor.

In this study, we have constructed a comprehensive map of cross-stress behavior by measuring the population fitness in pair-wise stress combinations for five stressors (osmotic, oxidative, alkaline, acidic, *n*-butanol) that are biotechnologically and biologically important ^3,9,35-37^. By performing genome-wide transcriptional profiling for each case, we identified key genes and pathways that are implicated in this phenomenon (**Figure 1A**). We then compared the observed cross-stress response arising from a short-term exposure to sequential combination of stresses to the response resulting from a longer term evolution of several *E. coli* lineages over 500 generations under the respective stressors. We identified the genetic basis of these differences by genome-wide re-sequencing of the mutants. Several hypotheses related to the stress order, stress combinations, mutation reversal and evolution in varying environments were tested through further evolving cells already adapted to *n*-butanol and osmotic to a total of 1000 generations under the respective conditions (**Figure 1B**). Fitness assessment and re-sequencing of the evolved strains provide insight on the genetic basis of acquired stress resistance and cross-behavior, while functional analysis highlight pathways that play a central role in this phenomenon.

**Figure 1.**
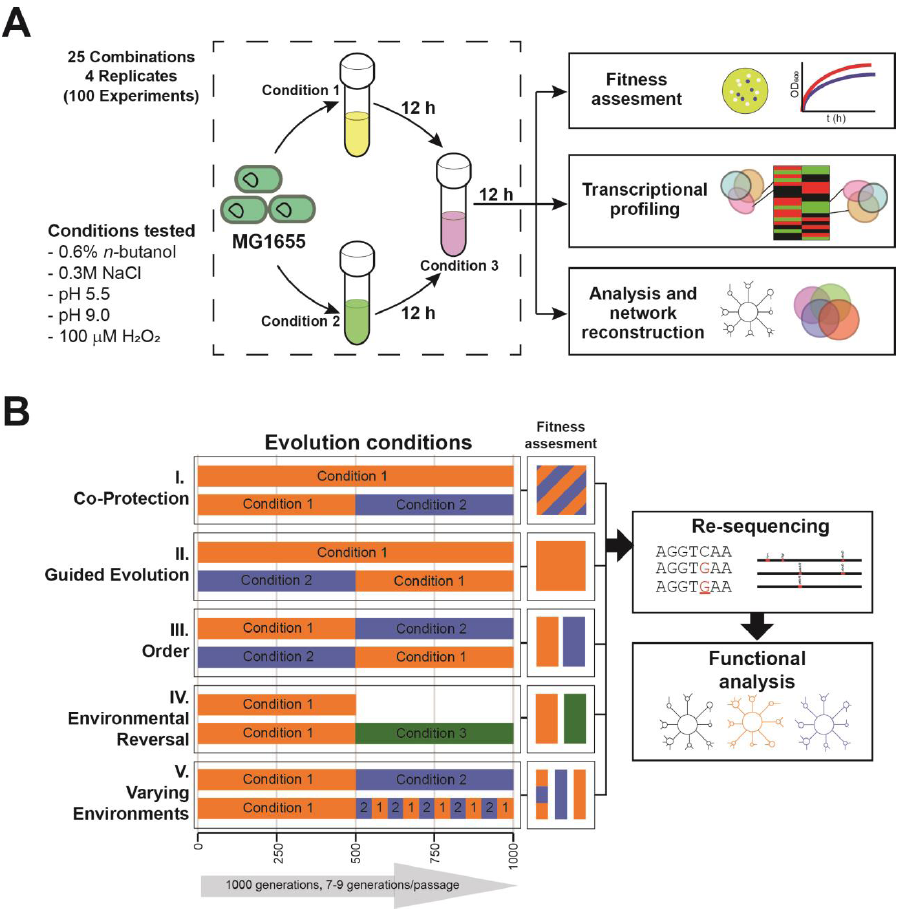
Overview of the experimental approach. **(A)** E. coli strains (4 biological, 2 technical replicates) were pre-conditioned in a first stressful (condition 1) or control (condition 2) environment for 12h before they were exposed to a second stress (condition 3) for an additional 12h. Competition assays, growth curves, genome-scale transcriptional profiling and functional network analysis was performed to assess the cross-stress behavior of E. coli for combinations of five stresses (n-butanol, osmotic, acidic, alkaline and oxidative stress). **(B)** Sequential evolution in stressful environments and hypotheses testing related to the adaptive potential of cross-stress behavior. E. coli populations were adapted over 1000 generations under various stresses to assess hypotheses related to co-protection, guided evolution, order, environmental reversal and varying environments. Genome-wide re-sequencing was used to map the genotypic landscape and phenotypic characterization was performed through competition assays and growth curves. Functional and pathway analysis was performed to identify the biological processes and network neighborhoods that are implicated in the acquired stress resistance

## Results

### Comprehensive map of the cross-stress behavior in *E. coli*

We tested the cross-stress behavior of *E. coli* MG1655 cells in all the possible pair combinations across five stressful environments (*n*-butanol, osmotic, acidic, alkaline and oxidative, four biological and two technical replicates). Out of the 25 possible combinations, 18 pairs (72%) have significant cross-stress behavior which is in accordance with the presence of a general stress response in *E. coli* (**Figure 2A, Supplementary Table S2, Supplementary File S1**). The highest cross-protection was observed when bacteria were exposed to acidic stress prior to *n*-butanol stress (DF = 1.15 ± 0.04, *p-value* < 1.46·10^-3^). We only detected one significant case of cross vulnerability, in the case of bacteria moved from oxidative to acid stress (DF = 0.97 ± 0.01, *p-value* < 1.91·10^-4^). We found a significant cross-protection effect between acidic media and high salt stress (DF = 1.10 ± 0.02, *p-value* < 5.17·10^-4^), which is a cross-stress behavior that previous studies found to be inconclusive in *E. coli* cells ^16,38^ but is known to occur in *S. typhimurium* ^18^. We also detected a cross-protection similar to that described for *S. typhimurium* and *Listeria monocytogenes* ^13,15^ between osmotic and oxidative stresses (DF = 1.14 ± 0.02, *p-value* < 6.41·10^-3^).

**Figure 2:**
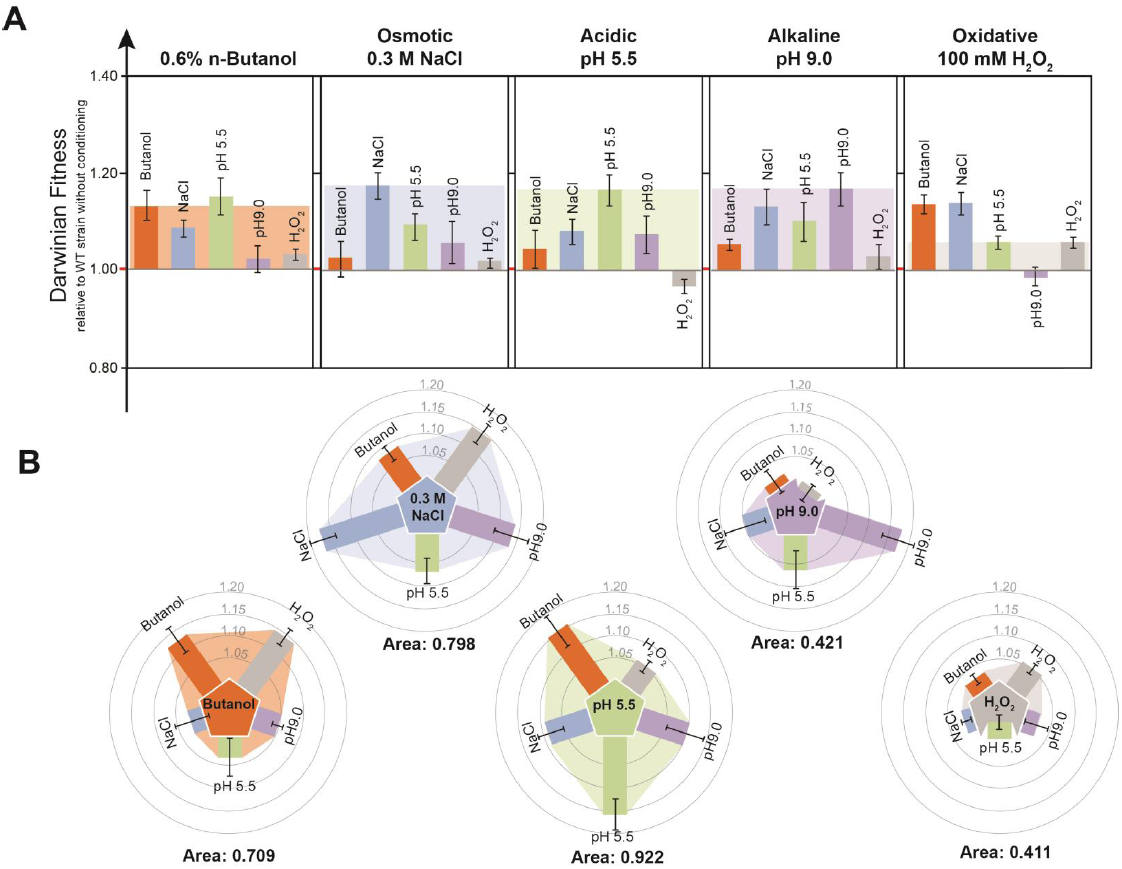
Hard-wired cross-stress behavior in *E. coli*. **(A)** Change in Darwinian fitness (W) of preconditioned E. coli populations (12h exposure) relative to the WT un-conditioned strain (W=1). Each sub-plot represents a different stressful environment where fitness was assessed through competition assays. Cross-stress protection ranges from 1.15 ± 0.04 to 1.02 ± 0.01 and cross-stress vulnerability is observed only in the case of E. coli strains pre-conditioned to oxidative stress before exposed to acidic stress (0.97 ± 0.01, p-value = 1.91 · 10^-4^). Shaded area depicts the fitness advantage when the population has been conditioned in the same stress. **(B)** Cross-stress area plots demonstrate the level of cross-stress protection for each conditioning environment.

A cross-protection was observed in bacteria adapted to either high salt concentration or *n*-butanol and then exposed to oxidative stress (DF = 1.14 ± 0.02, *p-value* <6.41·10^-3^ and DF =1.14 ± 0.02, *p-value* < 0.02, respectively). A recent study has identified cross-stress vulnerability for these two combinations ^39^, however in that case, *E. coli* was evolved for 500 generations in the first stress, hence accumulating mutations that can alter its cross-stress profile with respect to the ancestral line. Indeed, we found that evolved and un-evolved cell lines had significantly different behaviors in both the original growth media and the stress-induced M9 media supplemented with 100 mM H_2_O_2_ (**Supplementary Figures S1 and S2**).

In both presence of *n*-butanol and oxidative stresses, a first exposure to 4 out of the 5 stresses gives a selective advantage to the bacteria. Pre-adaptation to other stresses except acidic provides a fitness advantage in the oxidative environment, partially analogous to what has been described for *L. monocytogenes* ^19^. Surprisingly, in acidic stress we have observed the least cross-stress protection from all the other conditioned strains and we also observed the sole case of cross-stress vulnerability in the case of the oxidative-adapted strains (**Figure 2A**). On the other hand, bacteria adapted in low pH are well-positioned to compete in all other stresses. To quantify the degree of positive or negative cross-stress behavior, we introduce *cross-stress plots* (**Figure 2B**) where the acid-conditioned cells possess the highest area of cross-stress protection (0.922), while oxidative-conditioned cells have the lowest area (0.411) that also harbors the sole case of cross-stress vulnerability.

### Transcriptional profiles associated to cross-stress behavior

To analyze the underlying mechanism of cross-stress behavior we performed two types of transcriptional analysis. First, we performed a transcriptional profiling for each individual stressor at samples harvested 12 hours after exposure. Then we analyzed the transcriptional profiles after exposure to the second stressor (24 hours) to identify the differently expressed (DE) genes in the various cross-stress samples (**Supplementary Tables S3-5, Supplementary File S2**).

In the case of the genes differentially regulated in each stress, we identified 41, 203, 111 and 21 DE genes (*p-value* < 0.05 after Bonferroni correction) for *n*-butanol, osmotic, oxidative and acidic stress respectively. For each stress we merged results of the four different analysis (**Supplementary Figure S3**) and then data for each stress was used to make a new analysis that identifies DE found in more than one stress (**Figure 3A**). Interestingly, in acidic environments 18 of the 21 DEGs found (86%) overlap with other stresses that correlates well with the fact that this specific environment has the highest cross-stress area.

**Figure 3:**
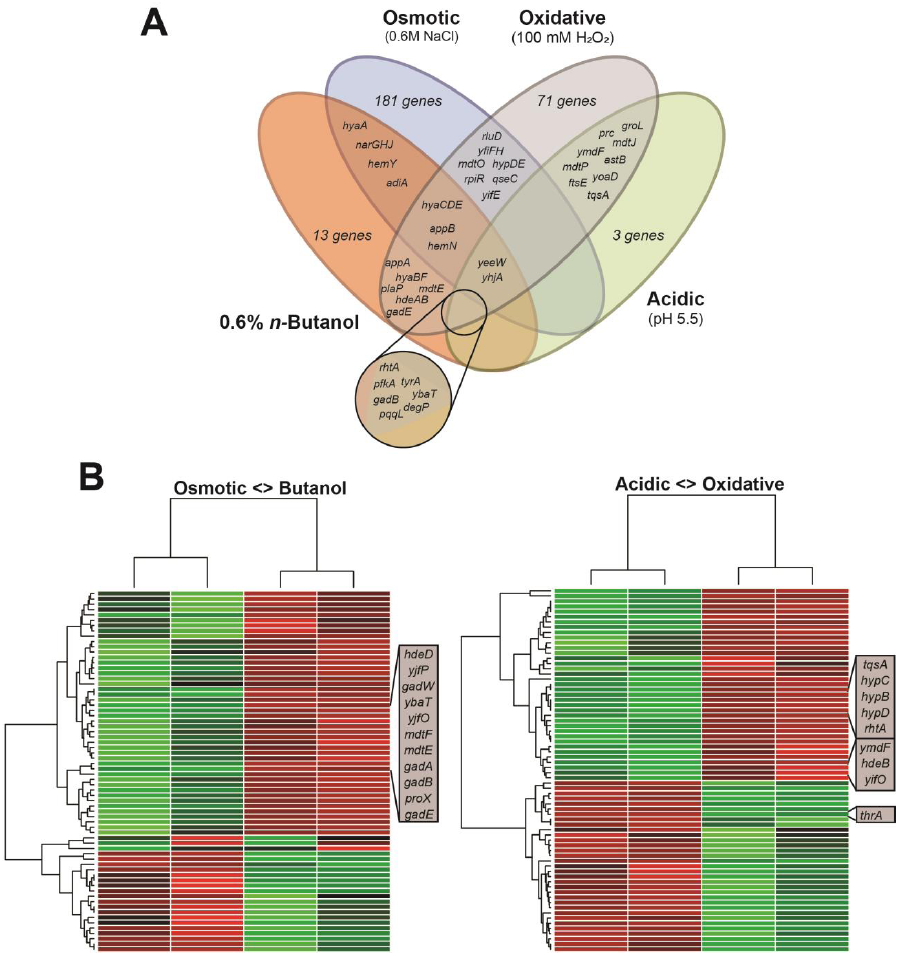
Transcriptional profiling in single stressors and cross-stress pairs. **(A)** genes differentially regulated in presence of one stress. **(B)** expression profiles of the cross-stress behavior. 4 analyses were perform, from n-butanol to osmotic, from osmotic to n-butanol, from oxidative to acidic and from acidic to oxidative. The expression profile was compare to a sample grown for 12 hours in M9 and then to the second stress (first column) and 24 hours in the second stress. Important implicated pathways are highlighted.

In osmotic and *n*-butanol stresses there is an under-expression of two operons, *nar* and *hya,* both reported to be over-expressed in anaerobiosis ^40,41^. The *hya* operon was also down-regulated under oxidative stress when compared to its expression under control conditions. The operon encodes for the synthesis of a hydrogenase that is implicated in pH stabilizing responses ^41^. The *gadE, gadB* genes from acidic resistance system AR2, the *adiA* gene from AR3 and the *hdeAB* acidic response ^42-45^ were found to be under-expressed under the osmotic and *n*-butanol stressor when compared to their expression in the absence of a stressor. As expected, the acidic resistance gene *gadB* is overexpressed under acidic stress conditions. We have identified both a common core of DEG for stress combinations and top ranked genes for individual stresses (**Supplementary Table S6, Supplementary File S2**). In osmotic stress, *proX* that belongs to the operon that encodes for osmoprotectant transportation is significantly overexpressed ^46,47^, while in oxidative stress we have high expression of the *yhjA* and *pfkA* genes. The first is a the *oxyR*-regulated peroxidase ^48^ while the second is of unknown function and we have recently showed that it is implicated with H_2_O_2_ sensitivity ^49^. Surprisingly, the highest ranked DEGs for *n*-butanol are the whole *hya* operon and the *gadB* gene, which are both related to acidic resistance. Indeed, culture in n-butanol was found to lower the pH to 5.9 after 12h, which explains the up-regulation of the acidic-stress cluster and the resulting cross-stress protection that was observed in this study.

To measure transcriptional changes in the profiles for both cross-stress protection and vulnerability cases, we performed RNA-Seq on four stress combinations: cells adapted to osmotic stress and then exposed to *n*-butanol (cross-stress protection) and cells adapted in oxidative stress that were subsequently exposed to acidic stress (cross-stress vulnerability) and their complementary pairs (from *n*-butanol to osmotic stress and from acidic to oxidative) to identify whether the observed difference in Darwinian Fitness can be attributed to the enactment of a dissimilar transcriptional program. Two controls were used as reference: cells grown for 12 hours in M9 and then 12 hours in the second stress and a population that has been exposed to the second stress continuously for 24 hours (**Figure 3B**). In all the analyses DEGs are similar when compared to both references, except from acidic to oxidative stress (**Supplementary File S2)**. When bacteria were exposed to *n*-butanol stress after being exposed to osmotic stress, there is only a small variation of expression compared to the bacteria that have been only exposed to *n*-butanol (9 DEGs only) showing that exposure to high salt conditions triggers a response similar to the response of bacteria exposed to *n*-butanol and no rewiring is needed giving the bacteria an advantage. Most of these 9 DEGs are acidic-response genes as found in the *n*-butanol samples like *gadB* and *gadE* (**Supplementary Table S7, Supplementary File S2)**. In contrast, when bacteria that has adapted in *n*-butanol face osmotic stress, more than 100 genes are differentially expressed when compared to the expression under osmotic stress exposure only. A similarly large differentially expressed gene pool is observed in the case of cells exposed to oxidative stress prior to acidic stress, which also is the only cross-vulnerability observed, where the list of DEGs goes over 750 genes, arguing for the cell’s need to enact a different expression program than the one used in the first stress in order to survive, which concords with a fitness disadvantage. In the 20 most significant DEGs (**Supplementary Table S8, Supplementary File S2**) there are several up-regulated genes previously described to be differentially expressed different stresses like acidic (*yjeI*, ^50^), manganese (*hflX*, ^51^), cobalt (*iscR*, ^52^) cadmium (*tyrA*, ^53^) and oxidative (*pfkA*, ^49^). In the inverse case, there is a cross-stress protection from acidic to oxidative and transcriptional profiling shows only 10 DEGs compared to the acidic grown bacteria that contains acidic related genes (*hdeB, gadB, hyaB*).

### Dissecting the memory imprint after adaptation to sequential stress combinations

Osmotic and *n*-butanol adapted cells demonstrate strong cross-stress protection both in the case of a short 12h exposure (**Figure 2A**) and adaptation after 500 generations ^39^. However, contrary to past results on adapted cells, where cross-protection has been found to be symmetrical, only bacteria exposed to *n*-butanol are protected from osmotic stress which argues for the existence of directionality of this phenomenon. To assess this hypothesis, we evolved osmotic and *n*-butanol adapted strains for an additional 500 generations in five independent experiments, all combinations of these environments, to further address questions related to cross-stress behavior after sequential stress adaptation (**Supplementary Table S9**). Growth rates of all the populations in media-only, osmotic stress and n-butanol showed that there was no growing defects after evolution (**Supplementary Figures S4-6, Supplementary Table S10, Supplementary File S3**).

The first hypothesis we tested is whether exposure to two stresses sequentially provides an advantage in an environment where both are present. We performed competition assays in an environment with both stresses (0.3mM NaCl and 0.6% *n*-butanol) between populations evolved for 1000 generations in a single stress (O1000, B1000) against those evolved in both stresses sequentially for 500 generations (O500B500, B500O500). Results show a significant fitness increase of the strains that have been exposed to both stresses (O500B500 *vs* B1000, DF = 1.16 ± 0.08; B500O500 *vs* O1000, DF = 1.24 ± 0.12) **(Figure 4A, Supplementary Table S11, Supplementary File S4)**. This *memory effect* is present long after the selection pressure for this particular stressor was removed (500 generations) and it can be attributed to the genetic background that has been acquired during the adaptation in the first stressor. In addition, this fitness increase is sufficiently large to overcome the additional fitness increase that is observed in the O1000/B1000 strains when compared to their O500/B500 counterparts due to their prolonged stress adaptation (DF = 1.12 ± 0.03, *p-value* < 7.15·10^-3^).

**Figure 4:**
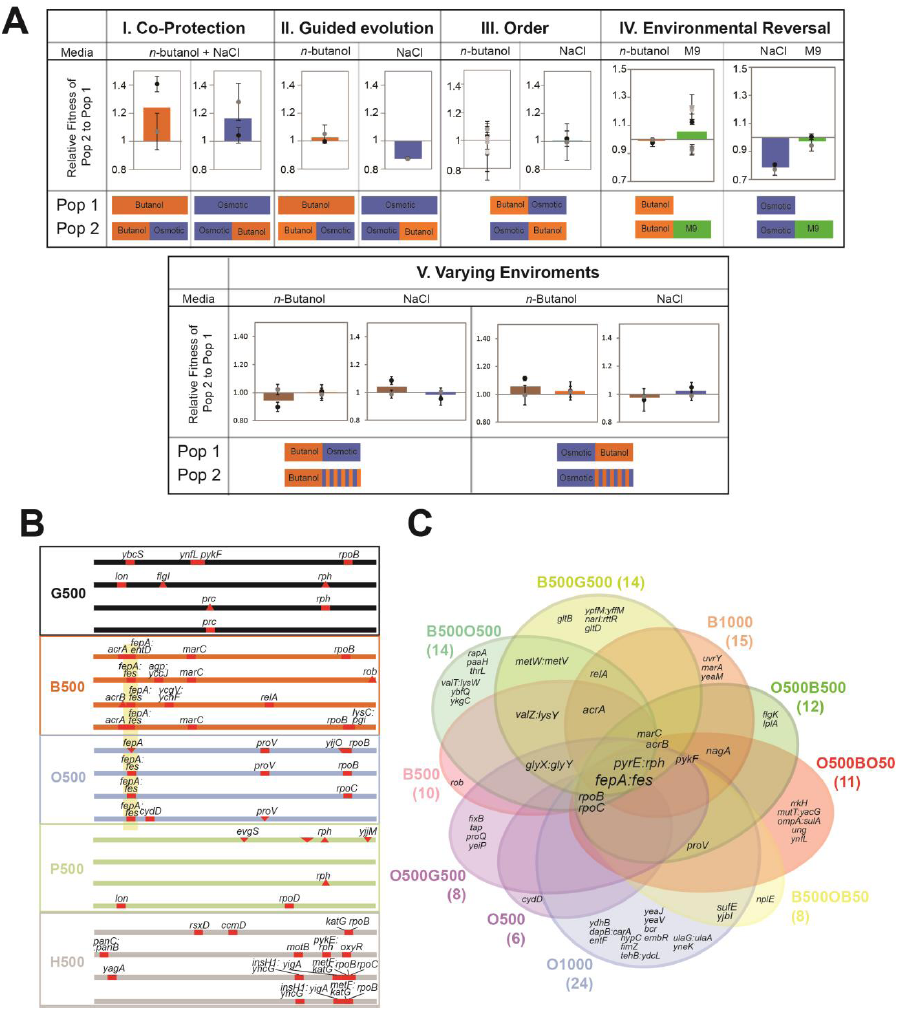
Phenotypic and genomic analysis of the populations evolved for 500 and 1000 generations. **(A)** Map of mutations in clones of populations evolved for 500 generations under one stress. Four clones of each stress were selected by fitness assessment. Mutations are draw relative to the reference genome of MG1655 and symbolized with a line if SNP and with a triangle if it is an insertion or deletion (upside and downside respectively). Details of the mutations can be find in Table S4. **(B)** Venn diagram illustrating the presence of the mutations in all the populations (Fig S6 and Table S10). **(C)** Results of the direct competition assays for each of the five hypothesis tested in this study. In all cases two biological replicates were performed (black and grey dots) but in the case of order in n-butanol and reversal of the n-butanol mutation in M9 were 6 replicates were performed.

Recent theoretical results show that the rate of evolution can be increased by guiding populations from states that are correlated and have increasing complexity ^54^. To challenge this notion in the case of osmotic and *n*-butanol adaptation, we competed populations evolved sequentially in both environments to those evolved in only one stress. In both cases the population evolved for a prolonged duration in a single stressor had similar or higher fitness than populations evolved in the two stressors sequentially (O500B500 vs. B1000, DF = 1.03 ± 0.02, *p-value* < 0.58; B500O500 vs. O1000, DF = 0.87 ± 0.01, *p-value* < 7.54·10^-11^). Concomitantly, we investigated if the order in which stressors are introduced play a role on the final fitness. We found that order does not have a significant impact to the fitness of the final populations when it is measured under competition in either stress (O500B500 vs. B500O500, DF = 0.99 ± 0.11, *p-value* < 0.99 in *n*-butanol stress; DF = 0.99 ± 0.010, *p-value* < 0.88 in osmotic stress).

We further probed on the environment’s impression on the genetic blueprint by investigating whether mutations are reversed when the stressor is removed. As such, populations that were adapted to environments for 500 generations with either *n*-butanol or osmotic stress (B500 and O500 populations, respectively) were competed to those where the initial 500 generations of stress-adaptation is followed by another 500 generations where the stressor is removed (medium-only environment; B500G500 and O500G500 cell lines). In the case of *n*-butanol, there is no significant difference in the Darwinian fitness when populations are competed under *n*-butanol stress (B500 vs. B500G500, DF = 0.99 ± 0.01, *p-value* < 0.34), while in the case of osmotic stress, the difference is profound in competition under osmotic stress, as beneficial mutations that are advantageous in osmotic stress have been lost (O500 vs. O500G500, DF = 0.79 ± 0.01, *p-value* = 8.30·10^-6^). In both cases, growth in media without the presence of stress was not affected (DF = 1.05 ± 0.04*, p-value* = 0.39; DF = 0.97 ± 0.03, *p-value* = 0.44).

Finally, to investigate the effect of alternating stresses to the evolutionary trajectory of each population, we evolved the O500 and B500 populations to alternating environments of osmotic and *n*-butanol stress (50 generations per environment, 10 switching events total; O500BO50 and B500OB50 cell lines). These populations have been competed with O500B500 and B500O500, respectively, in three distinct environments: *n*-butanol, osmotic stress and a varying environment with 8 hours for each stresses (**Figure 4A**). We did not observe any difference in the growth curves and competition assays between any of the populations grown in varying and static environments (B500OB50 and O500BO50 vs. B500 and O500 respectively; **Supplementary Table S11**).

### Acquired genetic mutations and their effect to phenotypic fitness during short-term evolution

Previous studies on the effect of single stress adaptation to cross-stress behavior have been focused on the identification of mutations on one cell line over short timeframes ^39^. To further elucidate the genetic basis of the acquired stress resistance and cross-stress behavior, we first sequenced four cell lines for each of the four of the stressors (osmotic, *n*-butanol, acidic and oxidative; alkaline stress was excluded because of media incompatibility) and the control environment (4 biological replicates, 5 environments, 20 lines total), where we have identified a number of loci that are mutation hotspots over two or more cell lines (**Figure 4B, Supplementary Table S12**).

Across all lines, two mutations were found to be synonymous, *ynfL* (G500) and *cydD* (O500) (**Supplementary Table S13**). The predominant mutations are in the intergenic zones *rph*:*pyre* (G500, P500, H500) and *fes*:*fepA* (O500, B500), as well as the sigma factors *rpoB* and *rpoC* (G500, O500, B500, H500). Interestingly, both *fes* and *fepA* are involved in siderophore enterobactin production that has recently been found to be linked with oxidative stress and M9 growth ^55^. Although its function is unknown, mutations in *rph*:*pyrE* have also appeared in a previous study in evolution on lactate ^56^. Both *rpoB* and *rpoC* encode for RNA polymerases and their mutations are very usual in evolution, more specifically mutations in *rpoB* have been linked to an increased evolvability and fitness ^57^ and mutations in *rpoC* to bacterial growth optimization in minimal media ^58^ and metabolic efficiency ^59^. We also found mutations in other genes already described like *pykF* in G500 which mutation is known to be beneficial for growth in M9 ^57^, *marC* and *acrAB* for resistance in *n*-butanol ^39,60,61^, *proV* in osmotic stress ^39,46,47^; *evgS* and *rpoD* in acidic stress ^39,42,62^ and *katG, oxyR, rsxD* and *ccmD* for resistance in oxidative stress ^39,63,64^. Interestingly, in oxidative stress we find mutations in the *yagA* and *yncG* genes, which is consistent with a previous finding that a deletion between *argF*-*lacZ* (that includes *yagA*) confers a high hydrogen peroxide resistance ^65^. In addition, *yncG* is homologous to Glutathione S-transferases known to help in the defense against oxidative stress ^66^. Other mutations with unclear importance in the stress where they appeared that were found are *ybcS*, *lon* and *prc* in G500; promoters of *yccF*, *ychF* and *pgi* and genes *rob* and *relA* in B500; *yijO* in 0500; *lon* in P500; and *motB*, *yigA* and the intergenic zone *panC*:*panB* H500. The mutations in the *ybcS, prc, yccF* and *yigA* genes are of unknown function, while others have known, but unrelated functions. For example*, lon* is a DNA-binding protease that degrades abnormal proteins ^67^, *ychF* is a ribosome binding catalase ^68^, *pgi* is an oxidative stress induced gene ^69^, *rob* is a transcriptional activator involved in antibiotic resistance ^70,71^, *relA* synthesizes ppGpp, an alarmone active under aminoacid starvation conditions ^72^ and *motB* is part of the flagellar motor ^73^.

We performed a similar re-sequencing analysis in the case of populations that have been evolved for a total of 1000 generations in a single or multiple sequential stress combinations (sequencing of multiple clones per population; **Supplementary Figure S7, Tables S14-16**). Four mutations were found to be synonymous and hence neutral, *paaH* (B500O500), *flgK* (O500B500), *fixB* in (O500G500) and *ynfL* (O500BO50). As shown in **Figure 4B**, the most common mutations are in the *fepA*:*fes* and *rph*:*pyrE* intergenic regions (8 and 7 cell lines, respectively) and mutations in the *rpoB* and *rpoC* genes (7 and 6 cell lines, respectively; **Table 1**). Many of these targets are mutation hotspots (**Supplementary Table S16)**, for example in the case of *rph*:*pyrE* and *fepA*:*fes*, parallel mutations have been detected in 9 positions that span 140 nucleotides (**Figure S8**) and 3 positions over 22 nucleotides (**Figure S9**) respectively. Populations of the B500 parental population contained several genes present in more than one population, *marC*, *acrA*, *acrB*, *relA*, *glyXVY*, with *valZ*:*lysY*. *marC* and *acrAB* have been previously described to be involved in *n*-butanol resistance ^39,60,61^ along with *nagA.* In contrast, populations evolved from the O500 background have only two common mutagenized loci present: *proV* described to be involved in osmotic resistance ^39,46,47^ and *nagA* present in osmo-tolerant cells ^35^.

**Table 1:**
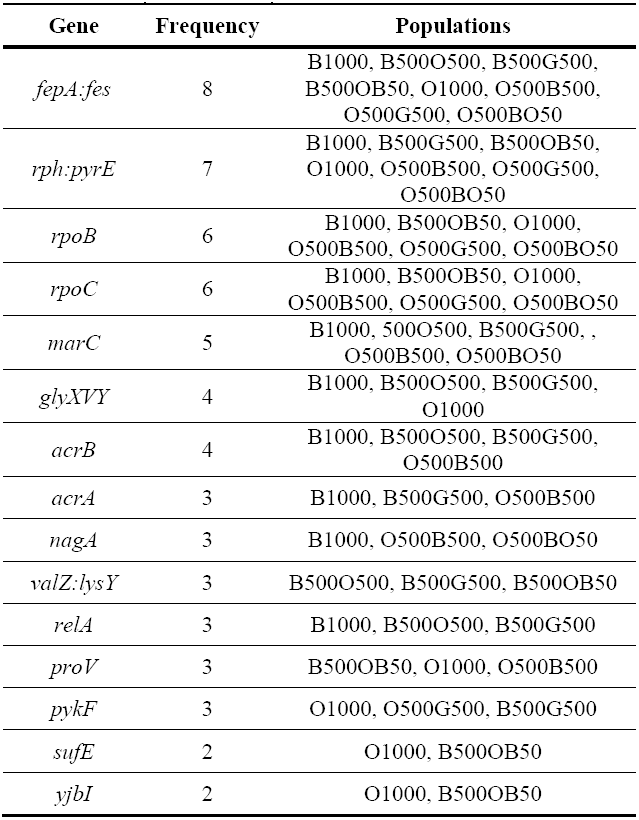
List of mutations that appear in 3 or more populations. Final count of populations is 8.

To decipher the genetic basis of this reversal, we compared the sequences of the B500 and O500 clones (**Figure 4A**) to those from B500G500 and O500G500 populations, respectively (**Supplementary Figure S7**). Cells that were first evolved in *n*-butanol and then adapted to stress-free media for 500 generations lost the *rpoB* mutation while acquired mutations in *pykF*, *gltD* and *gltB* genes. Similarly, cells that were first evolved in osmotic stress and then to stress-free media lost the mutation to the osmoprotectant gene *proV* and acquired mutations in the *pyrE*:*rph tap*, *yeiP* and *pykF* clusters. Although its function is unknown, mutations in *pykF* gives an advantage for growth in M9 ^57^. Both *gltD* and *gltB* encode genes for glutamate synthesis ^74^, *tap* produces a methyl-accepting protein ^75^ and *yeiP* is an elongation factor.

To associate genotype-to-phenotype, we performed growth experiments of single-knockout mutants ^76^ in medium-only, *n*-butanol and osmotic stress (**Supplementary Table S17, Supplementary Figure S10, Supplementary File S3**). Mutants in *pyrE* and *fes* cannot grow in any of the conditions tested, showing that the mutations found in their promoter affect the growth in M9 and are not involved in the stress adaptation. While mutation in *fepA* affects growth in all media, it provides a higher maximum growth rate in osmotic stress (**Supplementary Table S17**, media-only μ_max_ = 0.011 ± 0.000; *n*-butanol, μ_max_ = 0.014 ± 0.001; osmotic, μ_max_ = 0.020 ± 0.000). As previously described, Δ*pykF* enhances fitness in media-only ^57^, Δ*acrA* and Δ*acrB* in *n*-butanol ^39,60^ and Δ*proV* in osmotic stress ^46,47^. Other mutations that have a positive effect are Δ*relA* and Δ*sufE* in media-only, Δ*yjbI* in *n*-butanol and Δ*acrA* in osmotic. Previous results of repaired mutations under the mutant background elucidated the effect of *acrA* mutations in osmotic stress ^39^. Cross-stress trade-offs where found in the case of Δ*nagA* and Δ*sufE* that have a negative effect under osmotic stress. In constrast, no mutation was found to have a negative effect in growth under the absence of any stressor, or in presence of n-butanol.

### Functional and network analysis of cross-stress behavior

Re-sequencing and transcriptional profiling data were compiled in order to produce a network representation of stress resistance in *E. coli* (**Figure 5A, Supplementary Figure S11**). Network and functional analysis was also performed for each individual stress (**Supplementary Figures S12-S15**). Overall, enriched gene ontology clusters are related to motility, sulfur metabolic process, translation, DNA replication and cellular respiration, among others (**Figure 5A and B)**. Interestingly, the glycolysis/gluconeogenesis, response to drug genes and cellular respiration are involved in *n*-butanol response, with an *pykF* mutant from this pathway being unexpectedly fit, as it has the highest growth rate in these conditions. The importance of the *fes* and *fesA* genes of the Metal Iron Binding pathway is profound in the case of osmotic and oxidative stress and similar results have been obtained in the case of *fepA* iron transporter, in *n*-butanol, osmotic and acidic stress, highlighting the importance of metal transport in stress response. In osmotic stress, several translation and transcription pathways are found to be involved, while in acidic stress regulation of transcription and cell cycle clusters are enriched. The metal binding motif also includes a differential expressed gene, *yhjA,* that encodes for a cytochrome C peroxidase regulated by FNR and OxyR ^48^. This gene is been implicated in all stress responses and hence is an excellent target for further characterization. In this stress we also find motility genes involved in bacterial chemotaxis. In our analysis, a core network was identified with DEGs and mutations from every stress (33% *n*-butanol, 19% osmotic, 15% acidic and 33% oxidative). The fact that the members of the central clusters are all implicated in the acidic and osmotic stress correlates well with the previously shown *cross-stress plots* in which we showed that exposure to acidic and osmotic stresses have the higher cross-stress protection.

**Figure 5.**
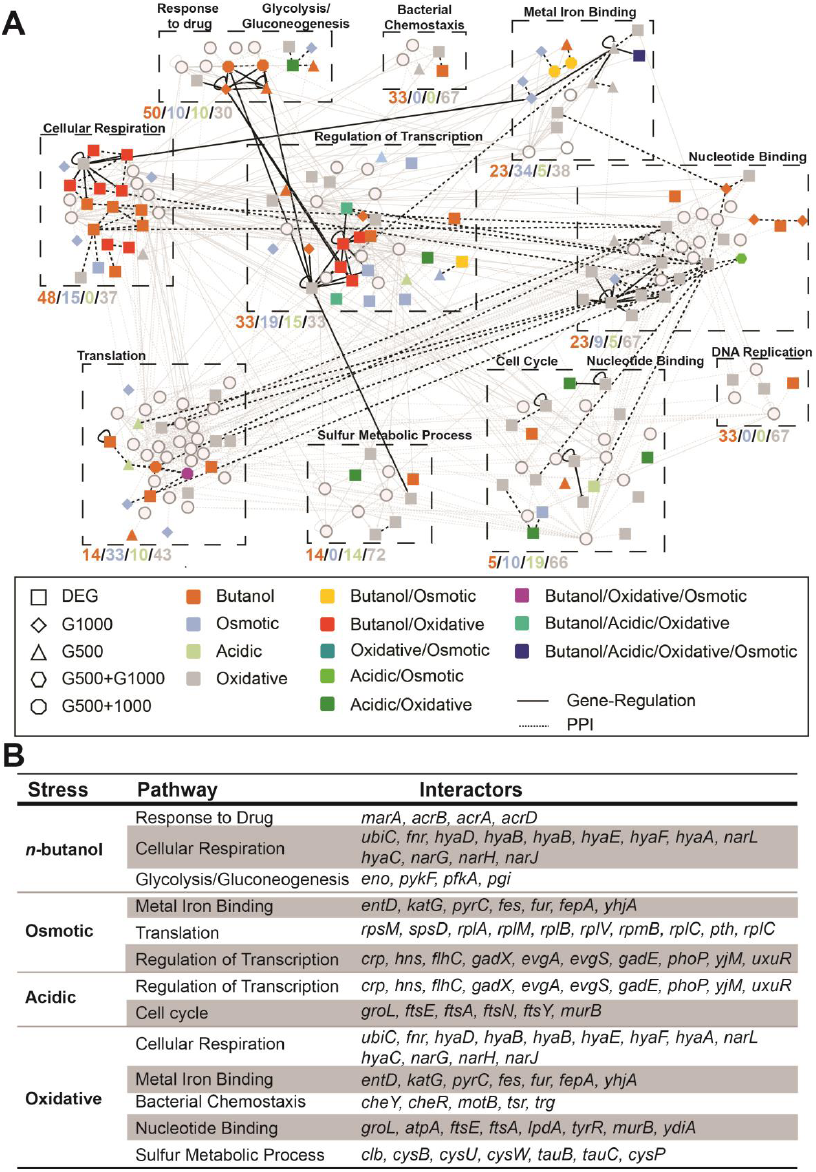
Network analysis and implicated pathways in stress resistance. **(A)** A functional network was constructed from PPI and TF-DNA data, superimposed with the re-sequencing and transcriptional profiling results of our analyses. Modularity-based algorithms were used to identify communities within the network, which were further analyzed for enriched clusters. The name of the most statistically significant cluster and the profile of the mutants/DEGs in terms of the corresponding stress (Butanol, Osmotic, Acidic, Oxidative, in that order) for each community is shown. For example, in the case of the first community, the Glycolysis GO term is the most over-represented and 50% of the observed mutations/DEGs in the community were identified in cell lines exposed/evolved in *n*-butanol. Light pink nodes are genes that are not mutated or DEGs, but connect two or more mutated/DEG genes in a path with a length shorter than three. **(B)** Highly enriched pathways and their members that are implicated in each stress.

## Discussion

A general observation that stems from our work is that exposure of a bacterial population to one stressor is generally beneficial, as 18 out of 25 cases tested in this work showed cross-stress protection with an average Darwinian Fitness of 1.08 ± 0.01 (*p-value* < 5.52·10^−8^). Interestingly, we also found the first case of cross-vulnerability when bacteria adapted to oxidative stress face low pH suggesting that some stress combinations can be used for a better sterilization. By performing transcriptional profiling we found that the number of differentially expressed genes across stresses is a good predictor of the underlying cross-stress behavior. Indeed, the case of cross vulnerability has the higher number of DEGs (>750), followed by the neutral cross-stress (>100) and finally both cases of cross-protection have a small amount of DEGs (9 and 10). Although the results here support this hypothesis, cross-stress protection can be also dependent on the DEGs of the first stressor as we find several genes with unknown or unrelated function, including the putative transcriptional repressor *rpiR,* the *yfi* operon and genes *pqqL* and *ybaT* that are present in all stresses but acidic (**Fig. 3, Supplementary file S2**). These genes constitute excellent targets for further experimentation to understand the mechanism under which they affect single stress and cross-stress behavior. Additionally, we identified several genes known to be differentially expressed under anaerobic conditions, which provide a clear link between genes induced by anaerobic respiration and stress resistance ^77^.

There is substantial similarly of the cross-stress behavior between *n*-butanol and osmotic stress, as well as a clear dissimilarity of these responses to that of oxidative stress with and without evolution. Analysis of the mutations found in the 4 clones sequenced (B500 and 0500, **Figure 4A**) discovered two genes in the *n*-butanol evolved populations that are also involved in oxidative stress that might be related with this differential behavior, the *ychF* catalase ^68^ and the *pgi* gene that are induced under oxidative conditions ^69^. Although not directly related to oxidative stress, in these clones we also found a mutation in the *rob* gene involved in antibiotic resistance ^70^. Unpredictably, in the O500 clones only one mutation is not related to the media, a mutation of the gene *yijO* which is of unknown function and remains to be investigated. Another explanation of the cross stress vulnerability is the *fepA*:*fes* mutation that, although it is expected to be related to the use of M9 as evolution media, it is only present in the B500 and O500 populations. The corresponding proteins belong to the enterobactin operon and have been recently implicated in growth under oxidative stress in M9 media ^55^ and thus its mutation might have a significant effect in M9 with oxidative stress.

If we are to predict where mutations will occur under evolution in a specific stress, it is reasonable to assume that the genes that are differentially expressed in that given stress will be likely candidates for polymorphisms or silencing. Comparison of the transcriptional profiles after short-term exposure and re-sequencing of evolved populations after 500 and 1000 generations argue that, in general, this hypothesis is not correct, as differential expression under exposure in a single stressor is not a proxy for mutation events during evolution in that stressor. Only in the case of osmotic stress we observe a mutation in the gene *proV* after evolution, which is in the same operon as the *proX* gene that was found to be over-expressed after the 12h exposure. Interestingly, after 1000 generations under *n*-butanol stress a mutation in *relA* appears and by network analysis we show that this gene is very close to several DEGs described in this stress like *gadE*, *mdtE* and *mdtF*. A similar case exists in oxidative stress where we detected a mutation in the intergenic region between *panB* and *panC,* with the network analysis showing that these genes have several interactions with other oxidative stress specific DEGs, such as *dnaJ*, *hypD* and *lolB*.

The evolved cell lines provide an interesting view of the evolutionary trajectories under the various stress combinations and it shows that their diversity is environment-specific. As shown in Figure 4B, cell populations that evolved under osmotic stress share few genetic mutations while populations evolved under *n*-butanol stress share many. Interestingly, despite the already described *n*-butanol related mutations, *acrA*, *acrB* and *marC*, we also found point mutations in several regions between tRNAs, *metW*:*metV*, *valZ*:*lysY*, *glyC*:*glyY* and *valT*:*lysW*. Mutations in four different spots argue against this being a random effect, which in turn implicates the tRNA production as a factor of *n*-butanol stress resistance. An expected result is the high abundance of mutations in the *fes*:*fepA* genes in all the populations as we already described this mutations in the parental population, B500 and O500. These mutations have been already described in similar scenarios and recently the involvement of the *fep* operon in growth in M9 ^39,55,56^. Surprisingly, another widely present mutation, *pyrE*:*rph,* is only present in the population evolved for 500 generations in a stress-free M9 medium, which argues for the *fes*:*fepA* mutation to be stress-related. We can track the emergence of the most common mutations by the number of polymorphisms detected in each mutation, for example the *fes*:*fepA* mutations appears in three different positions (**Supplementary Figure S9**) for the B500 and O500 populations. In the case of *rph*:*pyrE,* we have 9 distinct mutation positions (12 different mutations **Supplementary Figure S8**) arguing for the emergence of these mutations in the last 500 generations of the evolution. In addition, based on our results, we cannot conclusively dismiss or confirm the guided-evolution hypothesis although the O500B500 and B500O500 cell lines were found to have similar fitness to the 1000 generation populations, B1000 and O1000, in *n*-butanol and osmotic stresses, respectively (**Figure 4A**). This was surprising as the O500 and B500 have a significantly lower fitness with respect to their 1000 generation counterparts.

Finally, our last experiment involved fitness in a varying environment. More experimentation and in larger scale is needed to state with high confidence the magnitude and occurrence of this phenomenon. A recent work was also inconclusive regarding this phenomenon in *S. cerevisae* ^31^ with five times shorter cycles (approximately 10 generations per cycle). We also tested the hypothesis that shorter lag phases, instead of better fitness, should be observed under evolution in varying environments (New *et al*, 2014). Comparison of the μ_max_ of the varying environments population (**Supplementary Table S10**) shows that the O500BO50 general fitness is better than the O500B500 in osmotic stress (0.064 *vs.* 0.0525) but not in *n*-butanol (0.041 *vs*. 0.0375). In the case of the B500OB50 populations, a higher μ_max_ in osmotic stress than the B500O500 (0.0575 *vs*. 0.052) and a similar growth in *n*-butanol stress (0.039 *vs*. 0.043) was observed.

Although cross-stress behavior has been extensively studied in the past years, with many such cases related to food safety ^78^ and human health ^79^ this is one of the first studies where genome-wide profiling and a systems biology approach was used to create a comprehensive mapping of static and evolved cross-stress responses. This approach allows us to investigate the systemic response and molecular underpinnings of each condition combination, with the potential to inform new sterilization techniques and more efficient methods in clinical decision support ^80^.

## Acknowledgements

We would like to thank Athanasios Tsoukalas for his help with identifying the aminoacid effect of mutation events. This work was supported by NSF awards 1244626, 1254205 and DoD Army Research Office award W911NF1210231-0.

## Author Contributions

VZ performed cross-stress experiments, growth curves, strain characterization, and library preparation for genome sequencing and RNA-Seq. SQS performed the strain evolution and performed competition assays for validating the fitness of the populations. MK analyzed the sequences and the DEGs. NR performed the libraries for the X500 clones. VZ and IT evaluated data and prepared the manuscript. IT conceived of the study and supervised all aspects of the project.

## Materials and methods

### Bacterial strains and culture conditions

*E. coli* MG1655 and MG1655 Δ*lacZ* strains were used in all experiments of this study. The inclusion of Δ*lacZ* mutants allowed us to perform competition assays using X-Gal and IPTG staining. The neutrality of the Δ*lacZ* mutation was confirmed in all environments in a previous work of the laboratory ^39^. Minimal M9 salt medium with 0.4% (w/v) glucose as carbon source was used for cross-stress behavior competition assays. The stresses used are osmotic stress (0.3M NaCl), acidic stress (pH 5.5), alkaline stress (pH 9.0), oxidative stress (100 mM H_2_O_2_), *n*-butanol stress (0.6% *n*-butanol), and control (no-stress).

### Cross-stress behavior analysis

Two competing populations (*E. coli* MG1655 and *E. coli* MG1655 Δ*lacZ*) were pre-adapted to M9 overnight and inoculated at an approximate 1:50 ratio at a starting OD_600_ 0.004 in M9 with one stressor and only M9 for reference. After 12 h of growth, the same quantity of the strain adapted to stress and the strain grown in M9 were diluted in the second condition media (approx. 1:100 dilution) and competed for additionally 12 h. Samples were taken at 12 (time 0) and 24 hours (time 1). Four independent biological replicates each with two technical replicates were performed for each individual competition. Cell counts were determined on LB agar plates containing 0.25mM IPTG (Isopropyl β-D-1-thiogalactopyranoside) and 40 mg/ml X-gal (bromo-chloro-indolylgalactopyranoside). Plates were incubated at 37°C overnight. Darwinian Fitness (DF) was calculated as described in the Supplementary Methods section.

### Gene expression analyses

Cells were harvested at time 0 and 1. In all, 3ml aliquots were harvested and mixed with 1.5 ml 5% Phenol/ethanol (v/v) and stored at 80C until use. RNA was extracted using an RNeasy kit (Qiagen) and after first- and second strand cDNA synthesis, cDNA was broken using Diagenode Bioruptor NGS. End repairing, A tailing, linker ligation and PCR enrichment was made using the KAPA Library Preparation Kit (Kapa Biosystems). Size selection was performed with Agencourt AMPure XP (Beckman Coulter). After quality control, libraries were sequenced by Illumina HiSeq 2500.The low-quality raw reads were trimmed using Trimmomatic (v0.30) with default settings. Trimmed reads were aligned on most recent reference genome of *E. coli* MG1655 by using TopHat (v2.0.10) coupled with bowtie (v1.0.0) ^27,81^. The identification of the differentially expressed gene was done as explained in the Supplementary Methods.

### Adaptive Laboratory Evolution

Evolutionary adaptation experiments were performed in 10 ml volume in 125 ml shake flasks at 37C on an orbital shaker at 150 r.p.m. of an alternate of M9 with 85 mM NaCl (evolution media)media-only or with two stressors: osmotic stress (0.3M NaCl) and *n*-butanol stress (0.6% *n*-butanol). The OD_600_ of each culture was measured each day before the daily transfers to ensure that the estimated nine generations per day were reached (see Supplementary Methods).

### Whole-genome re-sequencing and mutation discovery of evolved strains

Selected clones were grown on LB medium overnight and genomic DNA (gDNA) was isolated using a Wizard Genomic DNA Purification Kit (Promega) and sequenced as described in the Supplementary Methods.

### Competition assays

Competing populations (*E. coli* MG1655 and *E. coli* MG1655 Δ*lacZ*) were grown under adaptive conditions overnight and inoculated at an approximate 1:1 ratio at a starting OD_600_ 0.004. Samples were taken at 0h and 24h. Two independent technical replicates were performed for each individual competition. Cell counts were determined as described before, using LB agar plates containing 0.25mM IPTG and 40 mg/ml X-. Plates were incubated at 37°C overnight. Darwinian fitness (W) was calculated as described in the Supplementary Methods section. In the case of the population adapted to a fluctuating media, the populations competed in a fluctuating media that varied from osmotic to *n*-butanol every 8 hours.

### Network analysis

We used TF-DNA binding data from RegulonDB ^82^and protein-protein interaction datasets ^83,84^ from the Bacterial Protein Interaction database ^25^ to construct a functional and regulatory network. We then mapped the results from our re-sequencing and transcriptional profiling analyses to build condition-specific sub-networks for each stress. Genes that connect either mutated or DE genes in paths with a path length of three or lower were also included in the network analysis. Modular organizations and community detection was performed on the resulting networks, through spin-glass model and simulated annealing techniques ^85^ using *igraph* R package ^81^.

